# A Compressive Sensing Approach for Inferring Cognitive Representations with Reverse Correlation

**DOI:** 10.1101/2021.09.02.458720

**Authors:** Benjamin W. Roop, Benjamin Parrell, Adam C. Lammert

## Abstract

Uncovering high-level cognitive representations of categories such as faces is an elusive goal that has been frequently sought using reverse correlation, a technique employed in fields ranging from neurophysiology to cognitive psychology. In reverse correlation, subjects are asked to make perceptual judgments (e.g., “do you see a face?”) about richly varying stimuli (e.g., white noise), and observed responses are then regressed against stimuli to yield reconstructions of the underlying cognitive representation. However, many thousands of stimulus-response pairs are frequently required, which severely limits the breadth of studies that are feasible using this powerful method. Techniques that are currently employed to improve efficiency, such as filtering the reconstruction, nevertheless bias the outcome. Here, we show that an advanced signal processing technique for improving sampling efficiency – compressive sensing – is directly compatible with reverse correlation. A trio of simulations are performed to demonstrate that compressive sensing can reduce the required stimulus-response pairs by up to 90% without biasing the reconstruction or can retrospectively improve the accuracy of the reconstructions on existing data. This work concludes by outlining the potential of compressive sensing to improve representation reconstruction throughout the field of neuroscience and beyond.

**Significance Statement:** Uncovering cognitive representations is an elusive goal that is increasingly pursued using the reverse correlation method, wherein human subjects make judgments about vague stimuli. Employing reverse correlation often entails collecting thousands of stimulus-response pairs, which severely limits the breadth of studies that are feasible using the method. Here we show that this methodological barrier can be overcome using compressive sensing, an advanced signal processing technique designed to improve sampling efficiency. Three sets of simulations are performed to demonstrate that compressive sensing can improve the accuracy of reconstructed cognitive representations and dramatically reduce the required number of stimulus-response pairs. This work concludes by outlining the potential of compressive sensing to improve representation reconstruction throughout the field of neuroscience and beyond.

## Main Text

Human perceptual experience is mediated by internal representations of the world around us. Representations bridge the gap between the raw information available from the senses and the abstract and often categorical nature of our perceptual experience by acting as reference patterns for everything from low-level spatial and temporal features (e.g., edges and shapes in the retina image (2)) to high-level characteristics of cognitive categories (e.g., faces (3)) or social constructs (e.g., the trustworthiness of a face (4)). Full understanding of perceptual experience therefore depends on the ability to characterize these internal representations, as they encapsulate important abstractions that are necessary to make sense of complex sensory inputs (Figure 1). While substantial progress has been made toward fully describing lower-level representations (e.g., visual and auditory receptive fields), similar characterization of higher-level representations has proven to be more elusive. This is largely because, despite their central importance to perceptual experience and behavior (5), such representations are notoriously difficult to measure (1, 3, 6-8).

**Figure 1.**
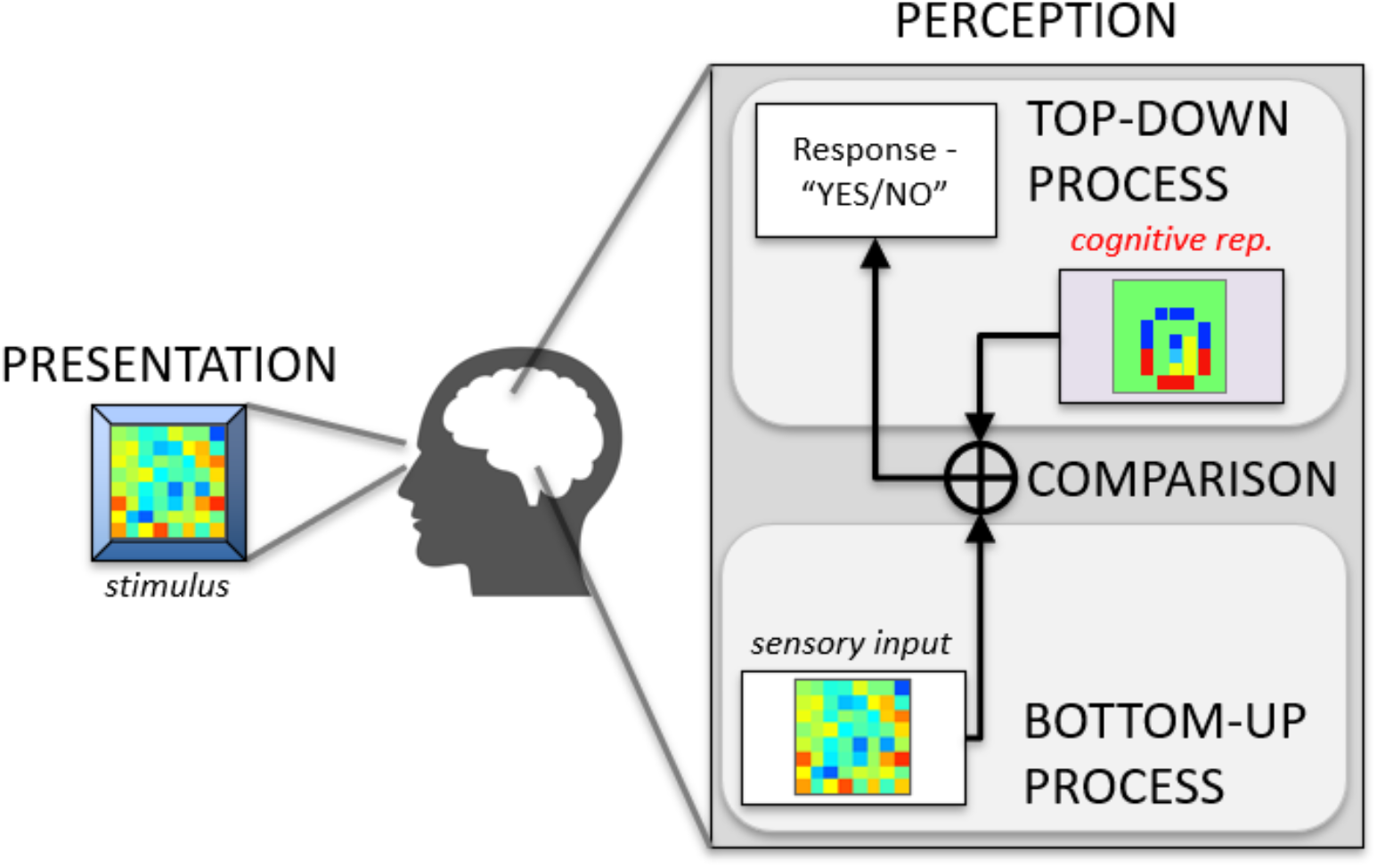
A schematic overview of perception. At its most fundamental level, perception is maintained by a complex cognitive system in the perceiver, involving the combined efforts of bottom-up and top-down processes that bridge the gap between sensory input and cognitive representations. Bottom-up processes extract relevant features from the raw sensory signal, while top-down processes relate incoming features to perceptual categories by comparing against prior expectations in the form of internal, cognitive representations. Such top-down theories are widely posited across multiple domains of human perception (e.g., (9)).

One of the more promising methods to overcome this difficulty attempts to uncover higher-level cognitive representations using a standard method for characterizing lower-level neural representations: reverse correlation (10). Reverse correlation reveals the latent representation in these low-level contexts by eliciting neural responses to richly varying stimuli (e.g., white noise) and regressing the observed responses against the stimuli over many trials. The same method can be utilized in the higher-level context by substituting voluntary responses for neural ones, in much the same way that one can report whether they “see” faces and other familiar shapes in clouds and other vague visual stimuli. Vague stimuli can thus be used to force the internal representations to exert their influence directly on elicited responses. In a pioneering study, Gosselin & Schyns sought to find representations of letters by asking participants that identify a letter “S” in visual noise stimuli (3); critically, the stimuli were purely white noise (and thus did not contain any actual letters), yet reverse correlation based on subject responses revealed a definitive “S” image recovered from each participant. Reverse correlation has subsequently been demonstrated to avoid the shortcomings of other techniques that bias expectations (11). Other groups have used reverse correlation to characterize the top-down processes of perception to abstract psychological categories, including the trustworthiness in voices (12), self-image (13), and “male” vs “female” faces (5) and shown that reverse correlation can quantitatively compare human decision-making to that of computers (14, 15).

Despite this initial success in certain contexts, reverse correlation has severe limitations that have prevented its wide-spread use for uncovering higher-level internal representations. Most critically, standard applications of reverse correlation require a burdensome number of stimulus-response samples for accurate reconstruction of cognitive representations with an adequate signal-to-noise ratio (8). For example, Gosselin & Schyns’ 2003 experiment required 20,000 trials from each participant; another required 11,400 (6). In cognitive and psychological experiments, the large number of trials combined with the long latency for measurable behavioral responses from subjects results in data collection protocols that can last weeks for each individual subject. This inefficiency limits the feasibility of applying reverse correlation to only those experimental protocols where subject participation and motivation can be maintained over long timelines. Even in such cases, the large amount of data required from each participant means the number of participants in a given study is typically very low (usually <5 (1, 3, 7)). This severely limits any insight into the generalizability of existing findings to a broader population.

Existing methods attempt to overcome this fundamental limitation of reverse correlation by biasing the input stimuli to decrease the number of required trials. This can be accomplished, for example, by generating stimuli through adding noise to known examples of some real-world category (e.g., an image of a face) rather than using pure noise, as done in (13). This increases the probability that subjects will report the stimulus as belonging to the category of interest but also biases the results toward representations consistent with the initial example. In order to realize the full impact of reverse correlation in uncovering cognitive representations, it is necessary to reduce the number of trials required without introducing such biases.

An alternative approach to improving the efficiency of reverse correlation is to impose constraints on the reconstruction process. Adding constraints can be a powerful way to improve reconstruction quality but can also severely limit the richness of the reconstruction in ways prespecified by the constraints themselves. For example, current approaches constrain reconstructions by smoothing with a 2D Gaussian kernel (1) or, more commonly, through low-pass filtering (e.g., (3)). While this can eliminate unwanted high-frequency noise from the reconstruction, it also presupposes that such high-frequency information does not form part of the underlying representation. For this reason, it is critical to assess the assumptions inherent in any added constraint and to be careful in deciding whether to utilize that constraint (16, 17).

One constraint that seems especially appropriate for reconstructing perceptual representations is that of sparsity. Sparsity is the notion that representations are composed of a finite and relatively small set of essential features. Importantly, sparsity is widely considered to be a fundamental principle of organization in perceptual systems at all levels (18), with strong empirical and theoretical support (18-21). Thus, sparsity is a fairly innocuous assumption that is highly likely to preserve the essential aspects and important variation in cognitive representations. This assumption can also be exploited to improve estimation of internal representations (17). Mineault (17) incorporated sparsity constraints in a binary logistic regression model to reconstruct visual-domain representations more efficiently, demonstrating an approximately 80% reduction in the number of trials to produce an equivalent quality reconstruction.

Recent advances in signal processing have resulted in a proliferation of efficient sampling methods based on the idea of sparsity, collectively known as *compressive sensing* (16, 17). To date, however, compressive sensing has not been applied to the reverse correlation paradigm. Here, we outline the underlying mathematical connections between reverse correlation and compressive sensing and provide a demonstration of the potential for dramatic efficiency improvements using the combined method. We will also show that, unlike previous approaches to incorporating sparsity constraints, it is possible to obtain reconstructions from compressive sensing analytically, using a closed-form solution. One such method by Zhang (22) is presented in detail and applied here in a novel way.

Compressive sensing begins with the assumption that signals of interest, x, which can include latent cognitive representations (Figure 2), are sparse (or, in the language of compressive sensing, *compressible*). This means specifically that they can be represented by a small number of functions from an appropriately-selected basis set (s *=* Ψ^T^_X_, for basis Ψ and weights s). If one assumes that responses, y, stem from a process of comparing stimuli to the latent representation (y=Φx, for stimuli Φ), it is possible to estimate the latent representation using only a small number of measurements by acquiring the basis function representation directly (i.e., y=ΦΨs) via sparse optimization approaches to find s. In practice, sparse representations can be found even when the chosen basis domain is quite general and incorporates no specific prior knowledge of the signal characteristics (e.g., the discrete cosine transform, wavelet transform).

**Figure 2.**
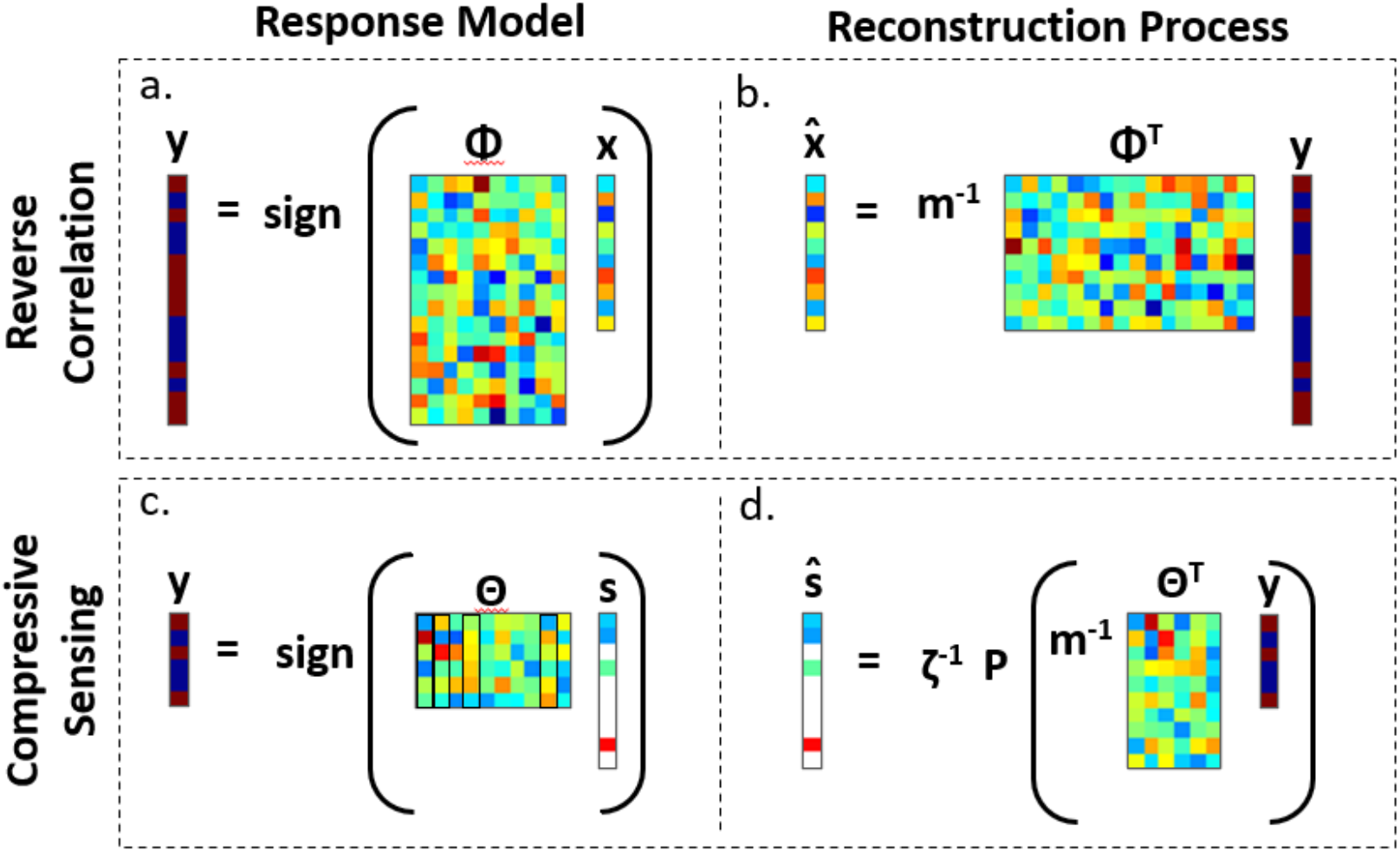
Process of reconstruction in reverse correlation and compressive sensing. (**A)** In reverse correlation, the vector of subject responses is modeled as resulting from the multiplication of a latent representation vector (x) and a stimulus matrix (Φ), which can be thought of as a similarity calculation between the latent representation and a vector representation of each presented stimulus. **(B)** An estimate of the latent representation 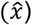 is then reconstructed by regressing responses against the stimuli. **(C)** In compressive sensing, the vector of subject responses is modeled as resulting from the multiplication of a sparse latent representation vector (s) and a compressive sensing matrix (Θ). The compressive sensing matrix is formed by multiplying a matrix of basis functions by the stimulus matrix (Θ=ΦΨ), which amounts to a similarity calculation between the stimuli and the known basis functions. **(D)** An estimate of the sparse latent representation (*ŝ*) is then reconstructed by regressing the responses against the compressive sensing matrix, soft-thresholding the resulting regression coefficients, and then normalizing by ζ=||P(m^-1^Θ^T^y)||,where m is the dimensionality of the stimulus vector and P is a soft thresholding function. The full representation estimate 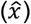 can be subsequently calculated using the estimated sparse representation and the known basis functions 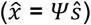 Note that the response vector y in compressive sensing is generally assumed to contain many fewer entries than in reverse correlation, and without sacrificing reconstruction accuracy.

Critically for the applicability of this method for reverse correlation, it has been demonstrated both theoretically (23, 24) and empirically (17) that using random (e.g., white noise) stimuli to elicit responses is a highly effective way to ensure accurate reconstruction of latent representations within the compressive sensing framework. Moreover, a significant portion of the substantial literature on compressive sensing has focused on the problem of inferring representations from binary responses (e.g., yes-no), a variation of classical compressive sensing called 1-bit compressive sensing (22, 25, 26). In other words, the somewhat unusual model of sampling assumed by compressive sensing maps directly onto the reverse correlation paradigm. This opens the door to potentially providing all the efficiency benefits of compressive sensing to the reverse correlation paradigm.

To directly assess the applicability of compressive sensing to recovering cognitive representations, we compare the results of simulations of Gosselin & Schyns’ original study on representations of “S” (Figure 3 top) using both conventional reverse correlation (3) and compressive sensing approaches. We used an image of the letter “S” as the target cognitive representation for reconstruction (Figure 3 top A). Specifically, the reshaped, vectorized “S” image was used as the representation vector, x (Figure 2), the basis for generating simulated subject responses in light of random stimuli contained in the stimulus matrix, Φ. The elements of Φ were generated by randomly sampling from a uniform distribution over the interval 0 and 1 and rounding those values to the nearest integer. A total of 10,000 subject responses were simulated using this procedure, representing the subject responses in (3). Additionally, a similar simulation was run using auditory spectrograms as stimuli. In both simulations, as in most cognitive experiments, we assume binary responses.

**Figure 3.**
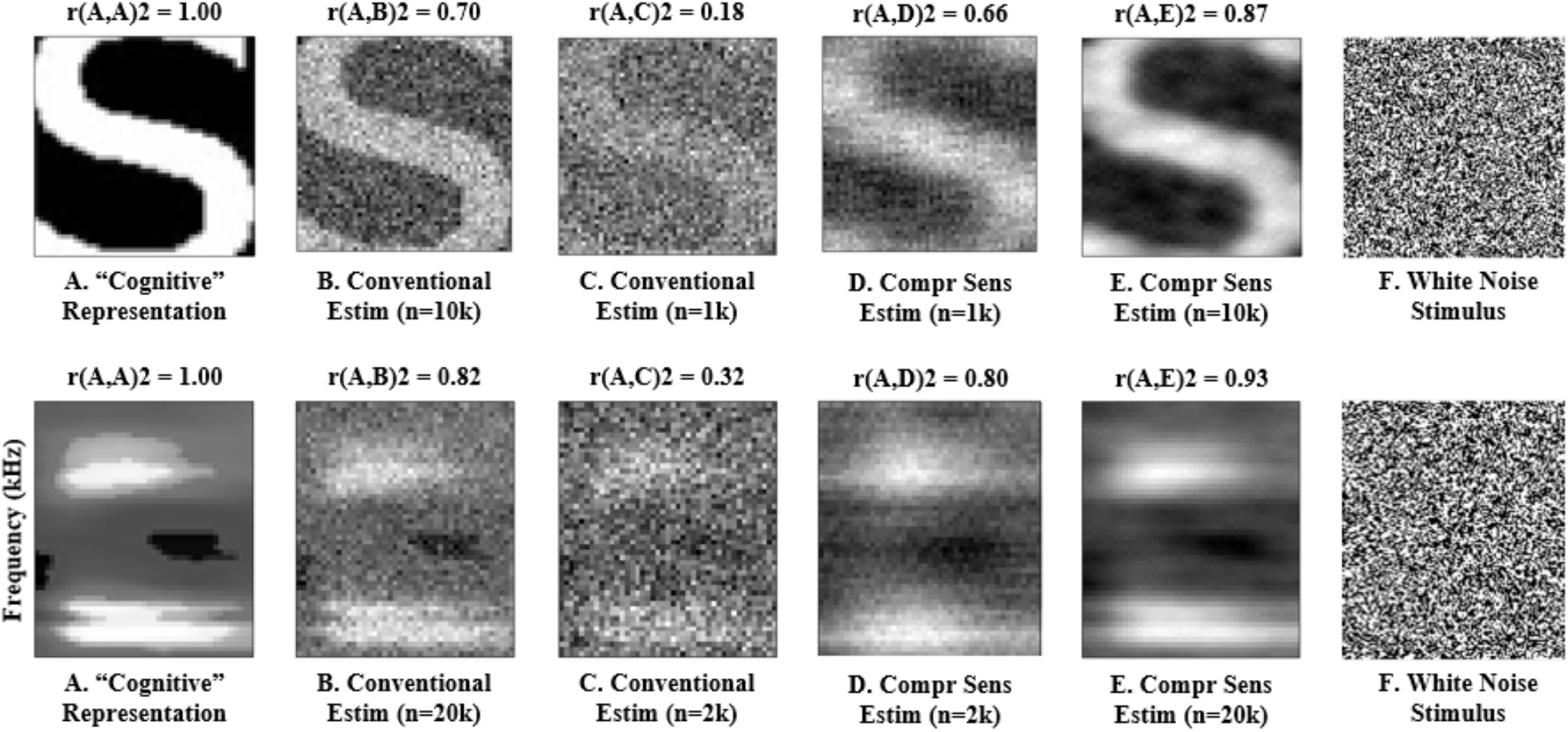
Comparison of conventional regression-based estimation and compressive sensing estimation. The template image **(A)** is estimated in **(B-E)**. An example white noise stimulus for each domain is shown in **(F)**. The method of estimation and number of samples used (n) is indicated below the image. The correlation coefficient (r^2^) between the template and the estimate is shown above the image, indicting estimation quality. Top row shows a simulation of the “S” from Gosselin & Schyns (3), and bottom row depicts a simulated time-frequency auditory spectrogram (an average phoneme spectrogram presented by Mesgarani et al. (27) is used as the template for reconstruction). In both simulations, compressive sensing provides equivalent accuracy as the conventional approach with only 10% as many trials.

The simulated subject responses were used as the basis for reconstructing the latent representation, x, using both conventional regression-based estimation and compressive sensing-based estimation. Reconstructions were generated using both the full set of stimulus-response pairs (10,000 or 20,000) and on a limited subset of a randomly-selected 10% of pairs (1,000 or 2,000). Estimation accuracy for a given reconstructed representation was assessed using 2-dimensional correlation coefficient (r^2^) between the target representation and the reconstructed estimate. Results (Figure 3) indicate that the image quality obtained using 10% of the samples via compressive sensing is effectively equivalent to that obtained using the full cohort of samples via conventional reconstruction, implying a 90% improvement in sampling efficiency. When allowed to operate on the full complement of samples, compressive sensing additionally shows improved accuracy over conventional reconstruction of approximately 15%.

The simulations reconstructing “S” and spectrograms showcase compressive sensing’s utility for analyzing novel data, yet compressive sensing is not limited to use solely in new studies. Rather, compressive sensing can also be retrospectively applied to existant data to improve results. To demonstrate this, further simulations were conducted to replicate the work of Smith, Gosselin, & Schyns, in which they used reverse correlation to reconstruct internal templates of faces (1). In that study, five subjects were each shown 10,500 white noise images and asked to identify those stimuli with a face in them; their results are recapitulated in Figure 4A. Compressive sensing simulations were implemented with the same number of samples as in the original work. To estimate the original stimulus-response data, a candidate measurement matrix, 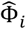, was randomly generated and used as the basis for estimating subject responses, *ŷ*, via the assumed data generating process 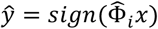, where *x* was the reconstructed face representation presented by (1). The estimated subject responses were then used to reconstruct an estimated face representation, which was subsequently subjected to smoothing using a Gaussian-shaped image filter as described by (1) to produce 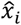. This procedure was repeated 1000 times, with the final measurement matrix and simulated subject responses being selected from the iteration which maximizes the image correlation between *x* and the estimate from that iteration, 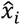. As may be expected in a sparse image, the compressive sensing reconstructions have a smoother, more face-like appearance (Fig 4B). Furthermore, Smith, Gosselin, & Schyns identified several facial structures in their reconstructions (e.g., a hairstyle on S3 and a nose, mouth, and chin outline for S1, S2, and S3), and these are more clearly observable in the reconstructions using compressive sensing. The reconstructions presented here are notably symmetric and lack high frequency variation. These properties are not enforced by the compressive sensing method. Rather, faces themselves are largely symmetric, and these reconstruction properties are driven by the data.

**Figure 4.**
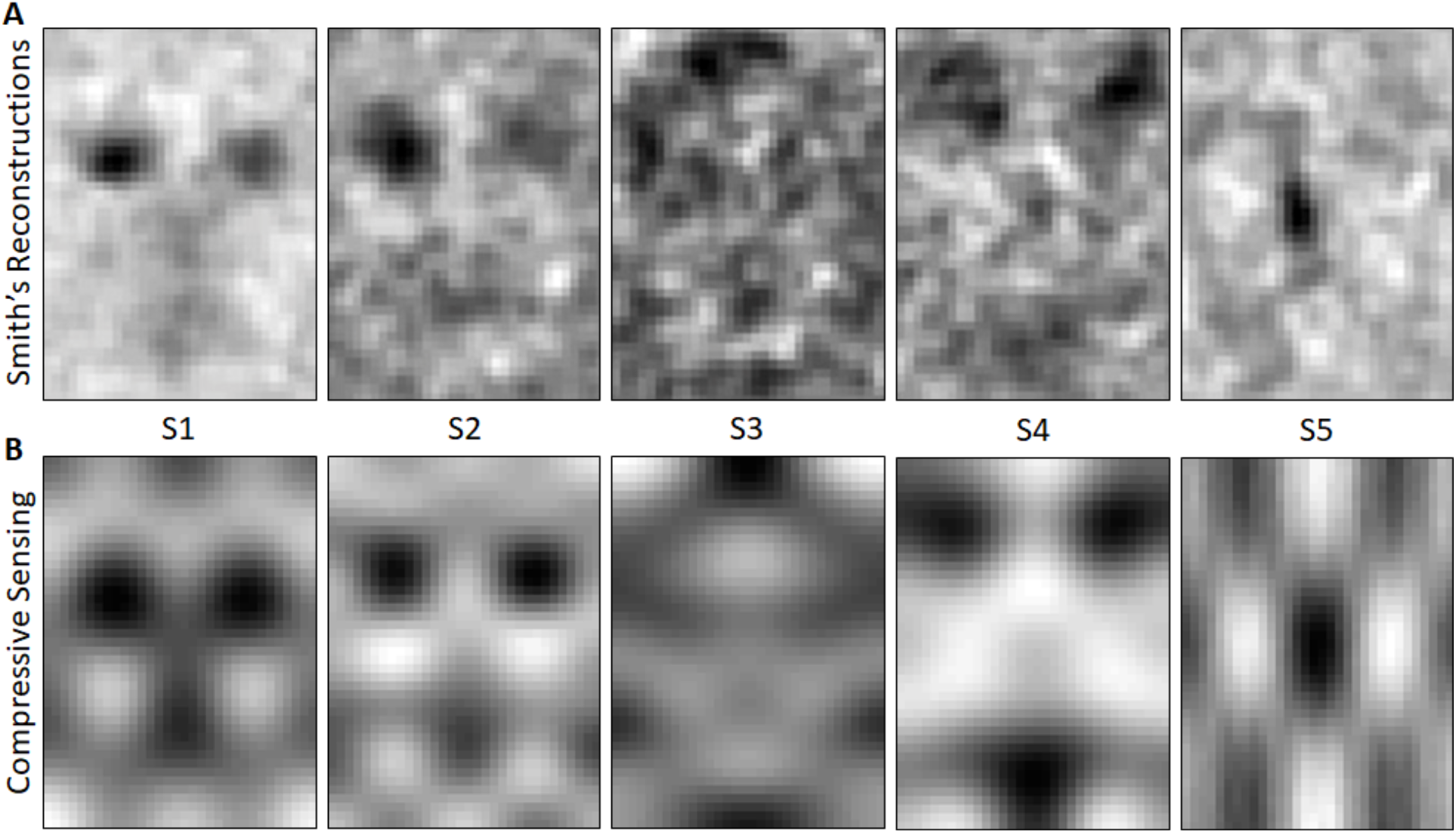
Comparison of conventional regression-based estimation and compressive sensing estimation of faces. **(A)** shows the reconstructions from (Smith, Gosselin, & Schyns 2012 (1)), and **(B)** depicts the compressive sensing reconstruction. Structural features identified in (A), such as a nose, mouth, and chin outline on S1, S2, and S3, are more noticeable in (B).

While validation on real data is necessary, these preliminary simulation results suggest that compressive sensing can reduce the number of trials required for accurate estimation of cognitive representations using reverse correlation by up to 90% without any loss of accuracy. Alternatively, more accurate reconstructions could be generated even while still reducing the number of trials, though by a smaller amount. Such a drastic increase in time- and cost-effectiveness would substantially increase reverse correlation’s potential for wide-spread use. Cognitive studies that would presently take weeks to complete could instead be performed in one sitting, and those studies that would otherwise impose limiting constraints on either the stimuli or the reconstructions (e.g., (13)) could be conducted with assumptions only about sparsity; this will lead to clearer, less biased representations. Outside of cognitive science, compressive sensing holds promise for drastically improving the efficiency and accuracy of analysis in any domain where reverse correlation is used, including a broad range of electrophysiology paradigms (e.g., (10, 28, 29)), calcium imaging (30), and auditory system function (31). That the reverse correlation responses can be continuous (e.g., neural firing rates (29)), ordinal (e.g., similarity scores (32)), or binary (e.g., yes/no (22, 25, 26)) gives compressive sensing widespread utility in areas of science even where reverse correlation is not currently in common use.

## Acknowledgments

We thank Karen Coghlan and Lori Ostapowicz-Critz of the Worcester Polytechnic Institute George C. Gordon Library.

